# Short-term memory capacity predicts willingness to expend cognitive effort for reward

**DOI:** 10.1101/2024.02.12.579951

**Authors:** Brandon J. Forys, Catharine A. Winstanley, Alan Kingstone, Rebecca M. Todd

## Abstract

We must often decide whether the effort required for a task is worth the reward. Past rodent work suggests that willingness to deploy cognitive effort can be driven by individual differences in perceived reward value, depression, or chronic stress. However, many factors driving cognitive effort deployment - such as short-term memory ability - cannot easily be captured in rodents. Furthermore, we do not fully understand how individual differences in short-term memory ability, depression, chronic stress, and reward anticipation impact cognitive effort deployment for reward. Here, we examined whether these factors predict cognitive effort deployment for higher reward in an online visual short-term memory task. Undergraduate participants were grouped into high and low effort groups (*n*_HighEffort_ = 348, *n*_LowEffort_ = 81; *n*_Female_ = 332, *n*_Male_ = 92, *M*_Age_ = 20.37, *Range*_Age_ = 16-42) based on decisions in this task. After completing a monetary incentive task to measure reward anticipation, participants completed short-term memory task trials where they could choose to encode either fewer (low effort/reward) or more (high effort/reward) squares before reporting whether or not the colour of a target square matched the square previously in that location. We found that only greater short-term memory ability predicted whether participants chose a much higher proportion of high vs. low effort trials. Drift diffusion modeling showed that high effort group participants were more biased than low effort group participants towards selecting high effort trials. Our findings highlight the role of individual differences in cognitive effort ability in explaining cognitive effort deployment choices.

**Significance statement:** We must often make decisions about when cognitive effort is worth the potential reward. Reward value, depression, and chronic stress in rodents can impact cognitive effort deployment decisions for reward, but factors like short-term memory ability can only be easily characterized in humans. We examined whether short-term memory ability, depression, chronic stress, and reward anticipation predict cognitive effort decisions for reward. In a short-term visual memory task with a choice of easier or harder trials for low vs. high reward, we found that only short-term memory ability predicted more choices of high vs. low effort trials. This research suggests that cognitive effort decisions could be driven by cognitive effort ability more than motivational factors like depression or chronic stress.

## Introduction

We are often faced with difficult choices about work and life. For example, we may choose to spend time thoroughly studying for an exam to gain a few extra percent points on a grade; alternatively, we may trade off this reward to work less and be able to do other tasks. These choices require trading off more or less effort for a larger or smaller reward, and thus involve deciding how much cognitive effort to deploy. In general, the motivation to deploy cognitive effort can be influenced by the potential reward to be gained (Shenhav, Botvinick, and Cohen, 2013; Shenhav, Cohen, and Botvinick, 2016; Yee, Crawford, Lamichhane, and Braver, 2021). Beyond this, an individual’s willingness to expend cognitive effort can also be linked to individual differences in factors that influence overall motivation, such as mood disorder levels (Grahek, Shenhav, Musslick, Krebs, and Koster, 2019; Pruessner, Barnow, Holt, Joormann, and Schulze, 2020; Yee, Adams, Beck, and Braver, 2019). Research in rodents has revealed patterns of individual differences in motivation to exert cognitive effort, where rats are classified as “workers” (high effort group) or “slackers” (low effort group) depending on whether they are willing to work more or less for reward, respectively (Hosking, Cocker, and Winstanley, 2016; Silveira, Wittekindt, Ebsary, and Winstanley, 2021; Silveira, Wittekindt, Mortazavi, Hathaway, and Winstanley, 2020). However, factors influencing individual differences in willingness to deploy cognitive effort for reward have yet to be fully characterized in humans.

In both rodents and humans, chronic stress (Birn, Roeber, and Pollak, 2017; Watt, Weber, Davies, and Forster, 2017) and depressive traits (Grahek, Shenhav, Musslick, Krebs, and Koster, 2019; Silveira, Wittekindt, Mortazavi, Hathaway, and Winstanley, 2020) have been shown to negatively impact cognitive effort deployment for reward. In rodents, chronic stress dampens reward anticipation even as acute stress heightens it (Ironside, Kumar, Kang, and Pizzagalli, 2018; Kúkel’ová, Bergamini, Sigrist, Seifritz, Hengerer, and Pryce, 2018). Here, chronic stress is typically induced by prolonged social defeat (Kúkel’ová, Bergamini, Sigrist, Seifritz, Hengerer, and Pryce, 2018) or non-social chronic mild stress, such as modifications to housing (Slattery and Cryan, 2017). In humans, chronic stress - unlike acute stress - cannot ethically be induced. Research into the impact of chronic stress on reward motivation primarily focuses on reports early childhood stress (Birn, Roeber, and Pollak, 2017; Watt, Weber, Davies, and Forster, 2017), and not shorter-term chronic stress of the kind that university students may experience (Towbes and Cohen, 1996). This latter form of chronic stress is more widely experienced than early childhood stress, and yet its impact on our willingness and ability to complete everyday cognitive tasks is not well understood.

Alongside chronic stress, depression is another motivational factor that has been shown to dampen reward anticipation (Terpstra, Vila-Rodriguez, LeMoult, Chakrabarty, Nair, Humaira, Gregory, and Todd, 2023; Westbrook, Bosch, Määttä, Hofmans, Papadopetraki, Cools, and Frank, 2020; Westbrook, Hankosky, Dwyer, and Gulley, 2018). In particular, anhedonia - a key symptom of depression impacting interpretations of reward (Slattery and Cryan, 2017) - can drive reduced reward anticipation and negatively impact willingness to deploy effort in rodents (Scheggi, De Montis, and Gambarana, 2018). Similarly, in humans, trait-level anhedonia (Treadway, Bossaller, Shelton, and Zald, 2012; Treadway, Buckholtz, Schwartzman, Lambert, and Zald, 2009) and anticipation of anhedonia (Sherdell, Waugh, and Gotlib, 2012) generally predicts reduced motivation to deploy physical effort for reward (Culbreth, Moran, and Barch, 2018). However, many tasks in life are cognitive as opposed to physical in nature; cognitive effort tasks may offer more naturalistic value as a result.

Decisions on when to deploy more or less cognitive effort require attentional control in both humans and rodents. This control can be driven by executive processes such as visuospatial working memory, or the ability to hold in mind and manipulate object locations in space. Increased working memory ability can, in turn, drive improved attentional control - an aspect of cognitive effort - while reduced working memory ability is associated with poorer control (Unsworth and Robison, 2020). Furthermore, during cognitively effortful tasks, reward anticipation modulates activation in neural circuitry that supports working memory (Fuentes-Claramonte, Ávila, Rodríguez-Pujadas, Ventura-Campos, Bustamante, Costumero, Rosell-Negre, and Barrós-Loscertales, 2015). Reductions in executive function capability - for example, given high levels of chronic stress and depressive traits - could drive excessive reliance on reward incentives, leading to maladaptive effort deployment that cannot be sustained (Kool, McGuire, Rosen, and Botvinick, 2010; Sandra and Otto, 2018). Although visuospatial working memory tasks can be conducted in rodents, including those exhibiting chronic stress and depression phenotypes, cognitive factors influencing willingness to deploy cognitive effort in humans are more complex than those that we can evaluate with rodents (Stephan, Volkmann, and Rossner, 2019). This is also just one of the many differences between humans and rodents in terms of how we use our sensory systems to navigate the world as well as in the reward cues for which we deploy effort; humans generally have a broader variety of primary and secondary rewards that they engage with in their everyday lives as compared to rodents. Acknowledging and accounting for these fundamental differences between human and rodent experiences of the world is an important yet underexamined element of translational research. Furthermore, humans require less training than rodents on cognitive effort tasks, and can be evaluated on a greater variety of behavioural measures revealing individual differences in willingness to deploy cognitive effort for reward. As such, it is important to translate rodent cognitive effort tasks into human tasks, allowing us to test human-based theories of cognitive and attentional control while maintaining comparability with rodent work.

In humans, the Expected Value of Control theory (Shenhav, Botvinick, and Cohen, 2013) predicts that, in general, people will deploy more cognitive effort when they know that they can effectively obtain greater rewards by doing so (Frömer, Lin, Dean Wolf, Inzlicht, and Shenhav, 2021). Additionally, the theory predicts that greater reward anticipation will lead to a higher expected value of deploying cognitive effort to obtain a reward (Grahek, Musslick, and Shenhav, 2020). Past work suggests that dopaminergic processes motivate physical and cognitive effort deployment decisions to maximize rewards (Michely, Viswanathan, Hauser, Delker, Dolan, and Grefkes, 2020) and to weigh rewards more heavily than effort costs. These processes also interplay with serotonergic processes that promote reward learning and biasing people towards more cognitively effortful options (Westbrook, Bosch, Määttä, Hofmans, Papadopetraki, Cools, and Frank, 2020) even when it is suboptimal to deploy effort earlier – which could occur more often in people with low levels of short term working memory ability (Raghunath, Fournier, and Kogan, 2021). However, the degree to which individual differences in reward anticipation and executive function capacity influence the likelihood of choosing to deploy more cognitive effort for higher reward is not known.

In rodents, different levels of motivation between ‘workers’ and ‘slackers’ have also been associated with differences in learning rate and bias towards choosing high effort trials, which may be linked to differences in cognitive capacity. Drift diffusion modeling in rodents completing a cognitive effort task has revealed that ‘workers’ with high accuracy may accumulate evidence more quickly towards selecting high effort, high reward trials than ‘slackers’ or ‘workers’ with low accuracy (Hales, Silveira, Calderhead, Mortazavi, Hathaway, and Winstanley, 2024). In an equivalent task in humans, such evidence could include awareness of one’s effort deployment willingness and ability as well as fatigue. For example, participants may build experience with how the task demands align with their short term memory capabilities, as well as their willingness to push these abilities further for higher reward. This evidence could also be used to make decisions about whether a higher reward is worth the additional effort. Overall, the above findings suggest that effort cost computations interact with reward anticipation as factors we rely on to make judgments about the value of deploying more or less cognitive effort for reward. In turn, individual differences in reward anticipation may be linked to depressive traits or chronic stress, and rodent research indicates that executive capacity may also influence choice behavior (Eichenbaum, 2017). The goal of the present study was to examine the degree to which individual differences in reward anticipation, chronic stress levels, and depressive traits on the one hand, and visuospatial short-term memory as an index of executive function on the other, predict human choice behaviour independently or in interaction.

### The present study

One way of examining factors that motivate humans to deploy more or less effort for reward is to offer participants the choice of completing an easier or harder cognitive effort task for a low or high reward, respectively (Treadway, Buckholtz, Schwartzman, Lambert, and Zald, 2009). In the current study, we built on an existing rodent choice task to investigate whether visual short-term memory capacity, chronic stress, depressive traits, and reward anticipation predict one’s choices in deploying cognitive effort for reward. A variety of human tasks exist to evaluate the costs of effort decisions. Some of these tasks examine the point at which high vs. low effort and reward options are equally preferred, including the Cognitive Effort Discounting task (COG-ED) (Crawford, Eisenstein, Peelle, and Braver, 2021; Westbrook, Kester, and Braver, 2013). In contrast, physical effort deployment tasks, such as the Effort Expenditure for Rewards Task (EEfRT) (Treadway, Buckholtz, Schwartzman, Lambert, and Zald, 2009), typically offer a binary effort choice structure with an easy trial for a lower reward or a harder trial for a higher reward. A key aim in our study was to examine whether human cognitive effort decisions can be modelled based on established animal models of cognitive effort deployment (Cocker, Hosking, Benoit, and Winstanley, 2012; Hosking, Cocker, and Winstanley, 2016; Silveira, Wittekindt, Ebsary, and Winstanley, 2021). Furthermore, the use of two effort levels encourages participants to strategize according to the trial difficulties available rather than fully discounting excessively easy or difficult trials offered in a parametric design. Thus, to ensure continuity with rodent cognitive effort tasks - including a binary choice paradigm - while maintaining a similar incentive structure to existing human cognitive effort tasks, we adapted the rodent Cognitive Effort Task (rCET) (Cocker, Hosking, Benoit, and Winstanley, 2012; Hosking, Cocker, and Winstanley, 2016; Silveira, Wittekindt, Mortazavi, Hathaway, and Winstanley, 2020) for use in humans. In the rodent paradigm, rodents first learn to associate illuminated lights with the opportunity to obtain a reward. In the main phase of the task, rodents are given the choice between a low effort, low reward (LR) trial or a high effort, high reward (HR) trial. Then, in a basic memory task, they must poke their noses in a hole that lights up for either 1000 ms (LR) or 200 ms (HR) (Cocker, Hosking, Benoit, and Winstanley, 2012). Rodents must rapidly encode the location of the illuminated light to succeed. Such a task can be scaled up to human working memory capabilities by including more stimuli and features that must be encoded to obtain a reward. For our study, we used harder and easier conditions (smaller vs. larger set size) from a visual short-term memory task (Luck and Vogel, 1997) to serve as high and low cognitive effort choices within the same kind of choice paradigm offered in the rCET. In each trial, participants could choose to perform an easier visual short-term memory task for a lower reward or a harder task for a higher reward. We examined overall visuospatial short-term memory capacity as well as indices of depressive traits (Beck, Steer, and Brown, 1996), chronic stress (Levenstein, Prantera, Varvo, Scribano, Berto, Luzi, and Andreoli, 1993), anhedonia (Rizvi, Quilty, Sproule, Cyriac, Michael Bagby, and Kennedy, 2015), and reward anticipation (Terpstra, Vila-Rodriguez, LeMoult, Chakrabarty, Nair, Humaira, Gregory, and Todd, 2023) as predictors of the tendency to choose lower or higher effort tasks. Furthermore, we used drift diffusion modeling (Hales, Silveira, Calderhead, Mortazavi, Hathaway, and Winstanley, 2024; Ratcliff, 1978) to examine factors that may contribute to choice biases.

## Materials and Methods

### Participants

We powered our study using a power analysis through the `pwrss` package (Bulus, 2023) to achieve an expected power of 80% at an odds ratio of 0.70 for a logistic regression evaluating the probability of a participant selecting 70% or more high effort trials (being in the high effort group in the study). This analysis gave us a target sample size of *N* = 487. We recruited *N* = 570 participants from the Human Subject Pool of psychology undergraduate students at the University of British Columbia; they were each compensated with bonus points for their courses as well as a CAD $5.00 Starbucks gift card. Of these, *n* = 42 participants did not complete the initial survey and *n* = 7 participants did not complete the reward anticipation task. Furthermore, *n* = 11 participants completed the change detection task more than once, *n* = 2 participants spent more than 30 seconds choosing any one trial - as this could indicate disengagement with task demands, and *n* = 79 participants performed at or below chance (50%) during the choice phase of the task. In total, we analyzed data from *N* = 429 participants (*n* = 92 male, *n* = 332 female, *n* = 5 other; Table 1). The study was approved by the Behavioural Research Ethics Board at the University of British Columbia, approval code H20-01388.

**Table 1.** Demographic information for all participants, by sex and effort deployment group. BDI prop = Beck Depression Inventory II proportion score (score divided by max score). PSS = Perceived Stress Scale score. DARS = Dimensional Anhedonia Rating Scale. Ant. = mean reward anticipation in the behavioural monetary incentive delay task. High effort group: > 70% HR trials selected; low effort group: <= 70% HR trials selected.

| Group | Sex | n | $M_{age}$ | $SD_{age}$ | $Min_{age}$ | $Max_{age}$ | $M_{BDIprop}$ | $SD_{BDIprop}$ | $M_{PSS}$ | $SD_{PSS}$ | $M_{DARS}$ | $SD_{DARS}$ | $M_{ant.}$ | $SD_{ant.}$ |
| --- | --- | --- | --- | --- | --- | --- | --- | --- | --- | --- | --- | --- | --- | --- |
| High effort | Female | 263 | 20.34 | 2.37 | 16 | 42 | 0.27 | 0.18 | 20.83 | 6.21 | 35.61 | 11.71 | 3.94 | 1.81 |
|  | Male | 80 | 20.70 | 2.34 | 17 | 32 | 0.19 | 0.14 | 17.69 | 5.30 | 37.69 | 9.62 | 4.29 | 1.75 |
|  | Other | 5 | 20.40 | 0.89 | 19 | 21 | 0.17 | 0.10 | 19.60 | 3.65 | 43.00 | 10.07 | 4.80 | 1.64 |
| Low effort | Female | 69 | 20.01 | 1.94 | 16 | 26 | 0.28 | 0.16 | 20.68 | 6.54 | 38.04 | 8.94 | 4.00 | 1.79 |
|  | Male | 12 | 20.92 | 2.97 | 18 | 28 | 0.19 | 0.14 | 21.08 | 4.38 | 39.17 | 8.61 | 3.96 | 1.51 |

### Stimulus Presentation

All participants completed the study online on their own devices, via the Pavlovia online study platform using PsychoPy 2021.2.3 (RRID: SCR_006571) (Peirce, Gray, Simpson, MacAskill, Höchenberger, Sogo, Kastman, and Lindeløv, 2019). Participants were not allowed to complete the study on mobile phones or tablets.

### Stimuli

All stimuli used in the study were generated by and implemented in PsychoPy (Peirce, Gray, Simpson, MacAskill, Höchenberger, Sogo, Kastman, and Lindeløv, 2019). In the first phase of the study, participants performed an online behavioural monetary incentive delay task as a measure of reward anticipation (Fig. 1A) as outlined in Terpstra, Vila-Rodriguez, LeMoult, Chakrabarty, Nair, Humaira, Gregory, and Todd (2023). In brief, during this initial task phase, participants saw a series of scales that they could click to indicate how excited they were to receive a reward. After a brief wait, participants would see a small happy face appear on one side of the screen. Afterwards, they were shown whether they received a reward or not, and were asked to view another scale where they would indicate how excited they felt. They completed this task for a series of eight trials.

**Figure 1.**
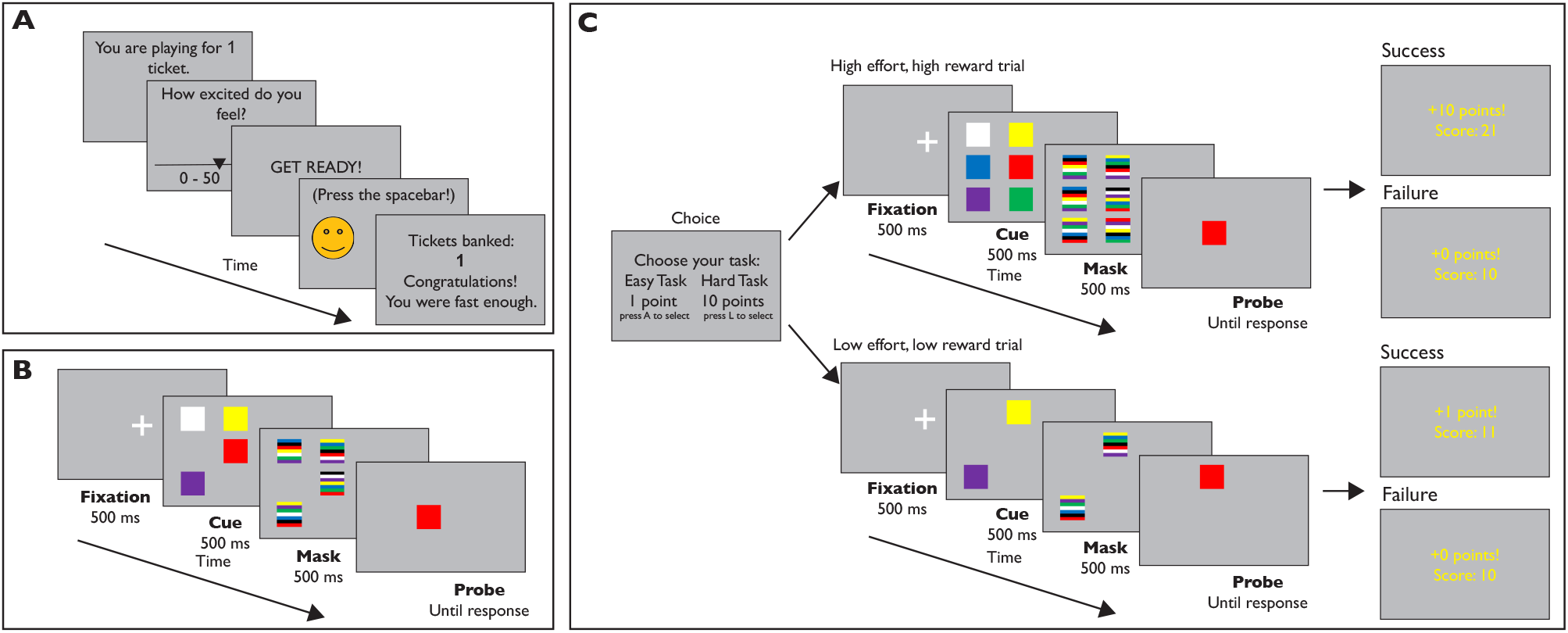
Layout of the experimental tasks. (A) The Monetary Incentive Delay Task, in which participants indicate their excitement in playing for tickets towards a monetary reward.(B) The calibration phase of the change detection task, where participants saw an array of either 2, 4, 6, or 8 squares and indicated whether the final square that appeared (the probe) was of the same or a different colour from the square that appeared in the same location in the previously show array. (C) The reward phase of the change detection task, where participants saw an array of either 2 or 6, or 4 or 8, shapes on screen depending on their performance in the calibration phase of the task. Once again, participants indicated with a keyboard press whether the final square that appeared (the probe) is the same or a different colour from the square that appeared in the same location in the array. Here, if they made the correct decision, they would receive 1 point (low effort trial) or 10 points (high effort trial); they would receive 0 points for an incorrect decision.

In the second phase of the study, as a measure of each individual’s visuospatial short-term memory capacity we presented a visuospatial short-term memory task (Fig. 1B, C) modified from Luck and Vogel (1997) and adapted from an open-source version of the task on Pavlovia (de Oliveira, João Roberto Ventura, 2023). In this task, we presented a series of trials with between 2 and 8 coloured squares that were presented for 500 ms. Each square subtended a visual angle of approximately 0.05° on the screen. A mask with multi-coloured squares would then appear at each of the original squares’ locations for another 500 ms, followed by a single coloured square appearing in one of the positions of the original squares. This final square had a 50% chance of being the same or a different colour from the square appearing in the same position in the initial part of the trial. After indicating with a keyboard press whether the square was the same or a different colour from the initial square, they would see whether or not they gained points towards a monetary reward.

### Procedure

#### Questionnaires and monetary incentive delay task

Participants began the study by completing an online questionnaire. After giving consent and demographic information, they were asked how about their history of depression and anxiety; COVID-related stress; the Perceived Stress Scale (PSS; Levenstein, Prantera, Varvo, Scribano, Berto, Luzi, and Andreoli (1993)); the Beck Depression Inventory II (BDI; Beck, Steer, and Brown (1996)); and the Dimensional Anhedonia Rating Scale (DARS; Rizvi, Quilty, Sproule, Cyriac, Michael Bagby, and Kennedy (2015)). Afterwards, they were redirected to the first phase of our study, the online behavioural monetary incentive delay task (Fig. 1A; Terpstra, Vila-Rodriguez, LeMoult, Chakrabarty, Nair, Humaira, Gregory, and Todd (2023)). This task measured participants’ reward anticipation via the mean of the excitement ratings given before each trial.

#### Visuospatial short-term memory task

Versions of this task served both as a means to measure individual differences in visuospatial short-term memory as well as the tasks of varied cognitive effort to be subsequently chosen for high/low effort trials. After completing the monetary incentive delay task, participants were redirected to the second phase of our study, the visuospatial short-term memory task (Luck and Vogel, 1997). Here, this task served as a means to measure individual differences in visuospatial short-term memory as an index of working memory. After receiving instructions on which stimuli would appear and how to respond to them, participants first completed a series of ten practice trials presented in a random order. On half of these trials, the probe square was the same colour as that of the cue square (congruent/change trial); on the other half, the probe square was a different colour from that of the cue square (incongruent/no change trial). Of these practice trials, four had a set size = 2, another four had a set size = 4, and another two had a set size = 6. In each trial, the probe square could be anywhere in the array, and participants were required to hold the whole array in visual short-term memory in order to successfully indicate whether the probe color had changed. Participants did not receive feedback on whether their responses were correct or not in the practice block, in keeping with Luck and Vogel (1997). Following these practice trials, participants completed (Fig. 1B) a series of 120 trials of the visuospatial short-term memory task with 60 change and 60 no change trials. In total, 30 trials of each set size (2, 4, 6, and 8 squares onscreen) were presented in a randomized order. Although the original task by Luck and Vogel (1997) contained trials with up to 10 squares onscreen, this largest set size was determined to be too difficult for participants to reliably complete correctly following piloting; as such, the maximum set size was 8. Furthermore, although the main task had a maximum set size of 8, the practice block only included a maximum set size of 6 so as to give a brief overview of the trials and required responses without giving excessive practice with a large set size, for which performance would be evaluated in the subsequent block.

Based on participants’ performance in this phase of the task, a K estimate of their visuospatial working memory capability (Rouder, Morey, Morey, and Cowan, 2011) was calculated for each set size, as follows:

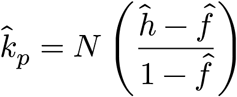

where *N* is the set size, 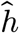 is the hit rate, and 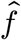 is the false alarm rate. As participants had to evaluate the whole display to judge the colour of the resultant single probe (Rouder, Morey, Morey, and Cowan, 2011), we used a whole-display K estimate measure (Pashler, 1988).

In addition to serving as indices of individual differences in visuospatial short-term memory, K estimate scores were used to calibrate the tasks used in the effort choice task (Fig. 1C), which was based on the rodent cognitive effort task (Hosking, Cocker, and Winstanley, 2016; Silveira, Wittekindt, Ebsary, and Winstanley, 2021). Here, participants completed a series of 30 trials. They would receive rewards for making a correct response, and they could choose a high effort, high reward (HR) trial or a low effort, low reward (LR) trial at each trial. Importantly in this task, trials with small and large arrays from the visuospatial short-term memory task served as the LR and HR trials. HR trials used a larger set size, but gave a reward of 10 points towards a monetary reward. LR trials had a smaller set size than the high effort/high reward trials, but yielded a reward of 1 point towards a monetary reward. We set the high effort trial reward to 10 points to motivate participants to continue deploying effort even for a much more difficult task. In order to ensure that the task difficulty in this phase was balanced by participants’ working memory capability, we used participants’ K estimate score at set size = 4 from the initially-presented visuospatial short-term memory task. This set size was determined through piloting to be the maximum set size before performance dropped off. Performance at set size = 4 was used to set a criterion for the available set sizes in HR and LR trials. Specifically, if the K estimate at set size 4 was <= 3, participants could choose an LR trial with a set size of 2 or an HR trial with a set size of 6. If the K estimate at set size 4 was > 3, participants could choose an LR trial with a set size of 4 or an HR trial with a set size of 8. Although participants were told that the the number of points they gained in this phase was proportion to the monetary reward they would receive, all participants received the same reward (a $5 CAD Starbucks gift card) at the end of the study. Afterwards, participants were redirected to a debriefing survey and received their course credit and monetary reward.

### Analyses

All analyses were conducted using R 4.3.1 “Beagle Scouts” (R Development Core Team, 2011) through RStudio (Booth et al., 2018).

The primary predictor variables in our study were 1) working memory ability, operationalized as a participant’s K estimate at a set size of 4; 2) depressive traits, operationalized as a participant’s BDI proportion score; 3) chronic stress levels, operationalized as a participant’s PSS score; and 4) reward anticipation, operationalized as a participant’s mean excitement before playing for tickets in the monetary incentive delay task (Terpstra, Vila-Rodriguez, LeMoult, Chakrabarty, Nair, Humaira, Gregory, and Todd, 2023). As our overall depression score measures were of more translational interest than anhedonia, given the role of depression in downweighing effort deployment and reward anticipation in both rodents (Slattery and Cryan, 2017) and humans (Grahek, Shenhav, Musslick, Krebs, and Koster, 2019), we focus on reporting this measure as opposed to the anhedonia measure we also collected. The primary dependent variable in our study was 1) proportion of HR trials chosen in the reward phase of the task; operationalized as whether participants chose the HR option for more or less than 70% of all trials in the reward phase. We further examined 2) accuracy, operationalized as the proportion of correct responses (hits and correct rejections) in the effort choice (reward) phase of the task and 3) choice latency, operationalized as the time in seconds until participants selected a difficulty level on each trial in the effort choice phase of the task. Finally, drift diffusion model parameters of drift rate, starting point, boundary separation, and non-decision time (Hales, Silveira, Calderhead, Mortazavi, Hathaway, and Winstanley, 2024) during choices in the effort choice (reward) phase of the task were additional outcome variables. To evaluate whether participants’ tendencies to choose high effort trials for high reward significantly explained performance (accuracy in the reward phase of the task) and choice latency (time until participants selected a difficulty level in the reward phase), we classified participants into two categories according to the criteria discussed in Silveira, Wittekindt, Ebsary, and Winstanley (2021): participants who chose the HR option for more than 70% of all trials in the reward phase were in the high effort preference group, while participants who chose the HR option for less than or equal to 70% of all trials in the reward phase were in the low effort preference group. This grouping was chosen to ensure continuity with comparable cognitive effort studies in rodents (Hales, Silveira, Calderhead, Mortazavi, Hathaway, and Winstanley, 2024; Hosking, Cocker, and Winstanley, 2016; Silveira, Wittekindt, Ebsary, and Winstanley, 2021). Although the grouping binarizes the outcomes of our analyses, it offers a clear point of translation from the rodent literature on which our task is based, and allows us to predict whether participants exhibit a high effort motivation vs. low effort motivation phenotype.

For our analyses, we first conducted a series of t-tests to evaluate sex differences in BDI and PSS scores. Past work suggests sex differences in depressive traits (Altemus, Sarvaiya, and Neill Epperson, 2014; Forys, Tomm, Stamboliyska, Terpstra, Clark, Chakrabarty, Floresco, and Todd, 2023) and chronic stress levels (Watt, Weber, Davies, and Forster, 2017), with women presenting with higher depression and chronic stress scores than men. For sex difference comparisons, only male and female participants were included as we were under-powered to report sex differences from those reporting sex as “other”. As participants may express high levels of both depressive traits and chronic stress - potentially impacting reward anticipation (Alloy, Olino, Freed, and Nusslock, 2016) - we also evaluated the extent to which BDI scores, PSS scores, and reward anticipation were correlated with each other. Next, we divided participants into high and low effort groups to ensure translational comparability with existing rodent work. As in Cocker, Hosking, Benoit, and Winstanley (2012), participants who chose the HR option for more than 70% of all trials in the reward phase of the task were in the high effort preference group, while those who chose the HR option for 70% or less of all trials in this phase were in the low effort preference group. We first conducted a binomial linear regression to determine whether sex, working memory ability, depressive traits, chronic stress levels, and reward anticipation significantly predicted whether a participant was in the high or low effort group in terms of their effort choices. As choices for high-effort trials may also vary continuously with visual short term memory ability, we then conducted a linear regression to evaluate whether the above regressors also predict the proportion of high effort trials selected. We then conducted two 2x2 within-between ANOVAs (Trial type x Group) using `anova_test` through the `rstatix` package (Kassambara, 2023) to evaluate whether accuracy or choice latency significantly differed by trial effort level (HR vs. LR) and motivation group (high effort vs. low effort groups). We examined accuracy on each trial and choice latency for selecting the trial difficulty to evaluate whether participants were matched for performance and time spent selecting a trial (Cocker, Hosking, Benoit, and Winstanley, 2012), regardless of how many high vs. low effort trials they selected. We then ran two multi-level models through the `lmerTest` package (Kuznetsova, Brockhoff, and Christensen, 2017) to evaluate whether sex, working memory ability, depressive traits, chronic stress levels, reward anticipation, and effort level significantly predict choice latency or accuracy.

Lastly, in order to evaluate whether the high and low effort groups differed in response strategies and biases towards selecting HR vs. LR trials, we fit a hierarchical Bayesian drift diffusion model, adapted from Ratcliff (1978) and run for 2000 iterations with 1000 warmup iterations and 4 Markov chains for Monte Carlo sampling, to data from all participants using the `hBayesDM` package in R (Ahn, Haines, and Zhang, 2017). We used this model to compare drift rate, starting point, boundary separation, and non-decision time between participants who chose high levels of effort vs. those who chose low levels of effort. The drift rate captures the rate at which participants drift towards a decision boundary (selecting an HR or LR trial). The starting point is expressed as a percentage of the distance between the lower and upper boundaries; it captures the initial bias, at the beginning of each trial, that participants have towards selecting an HR over an LR trial. The boundary separation, expressed relative to 0, captures a trade off between speed and selecting the HR trial as opposed to the LR trial. Lastly, non-decision time captures the part of choice latency that is not related to cognitive effort decisions, such as the time required to execute a motor response upon making a decision.

## Results

### Depressive traits and chronic stress levels

Summary statistics for depressive traits (BDI) and chronic stress levels (PSS) can be found in Table 1. After correcting for multiple comparisons using the Bonferroni method, women had significantly higher depressive trait scores (*t*(261.23) = 5.35, *p* = <0.001, *d* = 0.45) (Fig. 2A) and chronic stress scores (*t*(270.03) = 4.93, *p* <0.001, *d* = 0.40) than men (Fig. 2B). However, men and women did not significantly differ on anhedonia scores (*t*(253.79) = -2.05, *p* = 0.373, *d* = -0.17) (Fig. 2C) or levels of mean reward anticipation (*t*(233.53) = -2.50, *p* = 0.118, *d* = -0.23) (Fig. 2D). Levels of depressive traits (BDI), chronic stress (PSS), and reward anticipation were significantly correlated with each other; neither measure was significantly correlated with working memory ability (K estimate) (Fig. 3).

**Figure 2.**
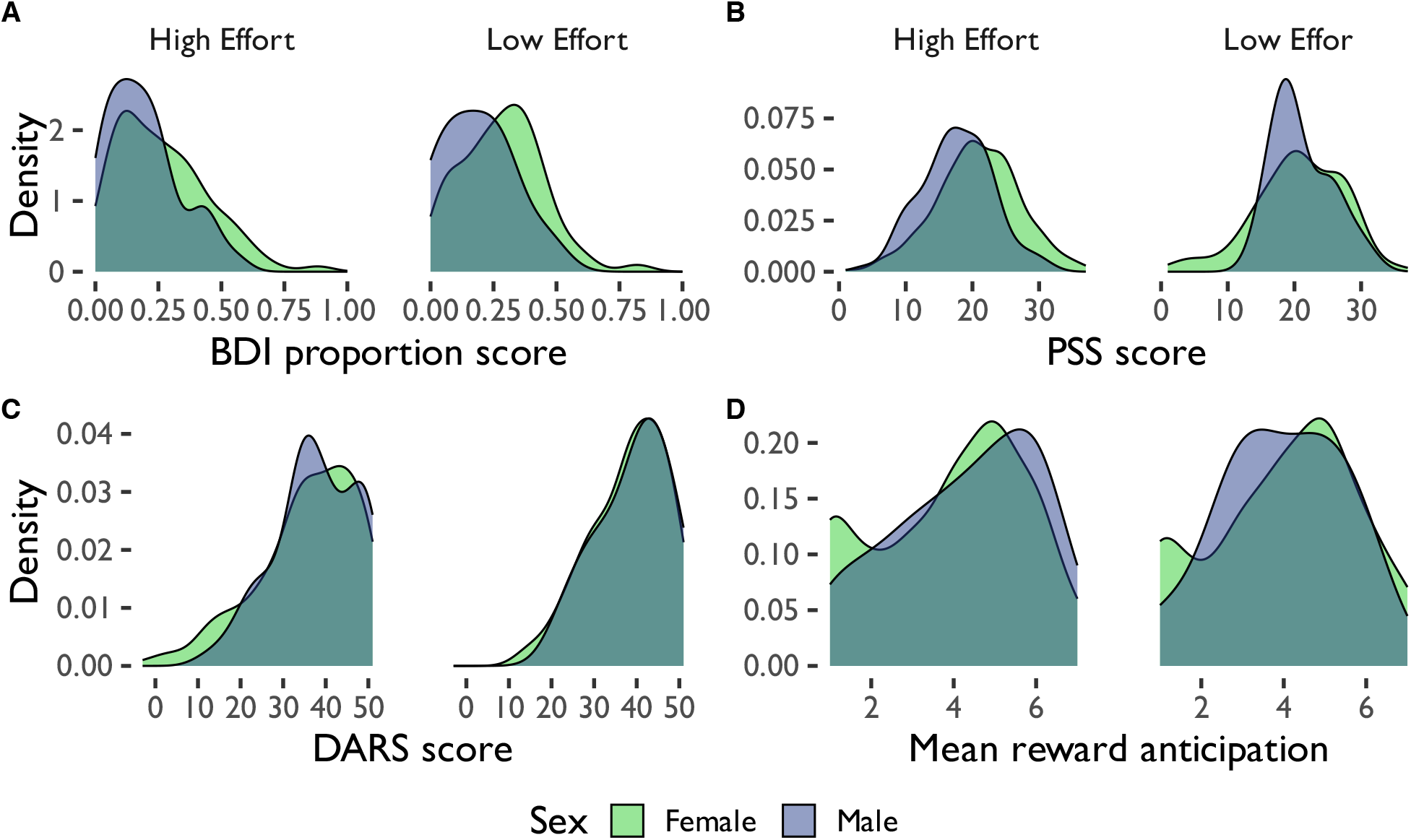
Distribution of (A) depressive trait (BDI) proportion scores (score divided by total possible score), (B) perceived stress (PSS) scores, (C) anhedonia (DARS) scores, and (D) mean reward anticipation scores by sex and effort deployment group. BDI = Beck Depression Inventory, PSS = Perceived Stress Scale, DARS = Dimensional Anhedonia Rating Scale.

**Figure 3.**
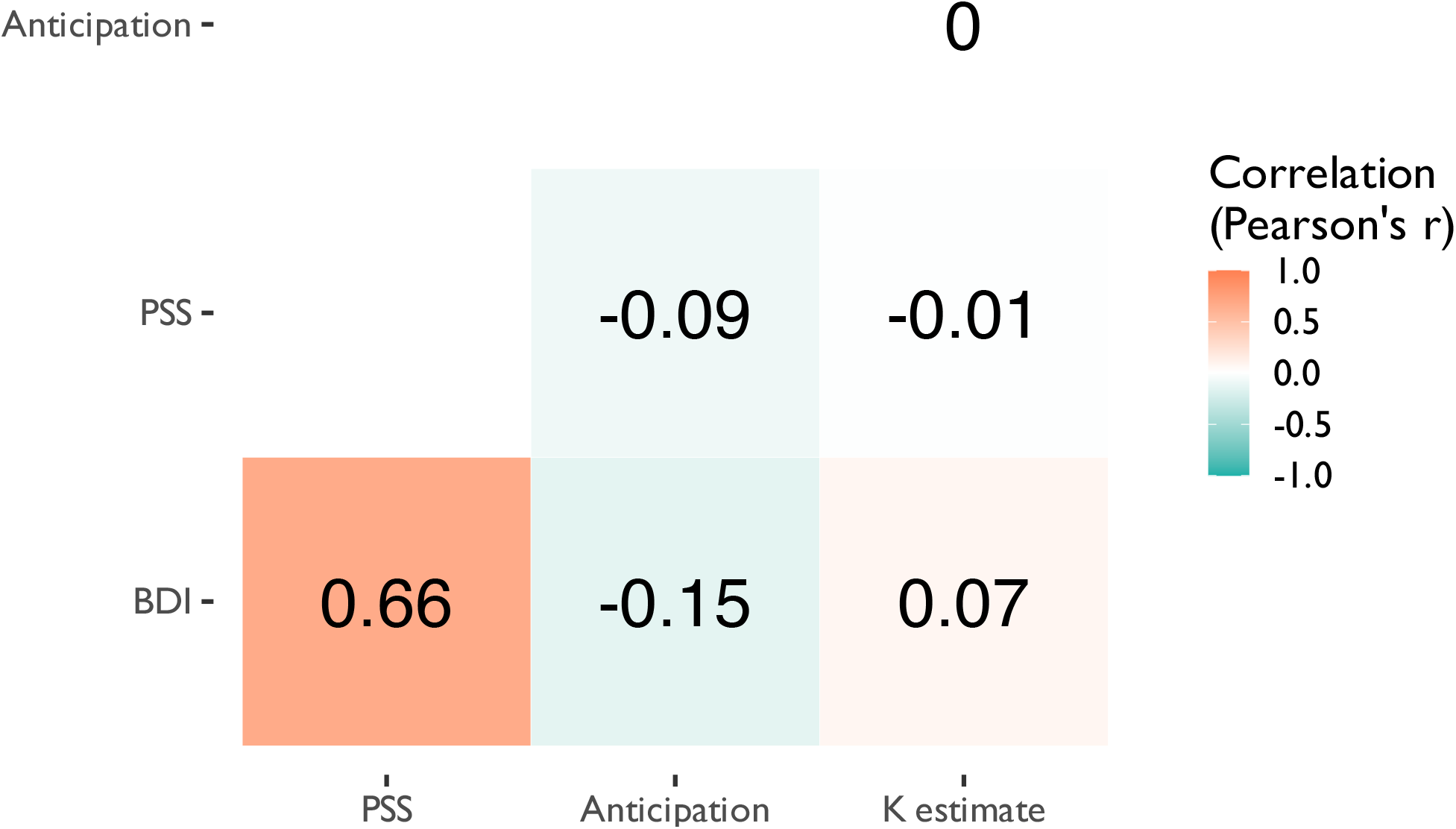
Correlations among depressive traits (BDI), perceived stress (PSS), reward anticipation, and short term memory ability (K estimate). BDI = Beck Depression Inventory, PSS = Perceived Stress Scale. Anticipation = mean reward anticipation. K estimate = visual short term memory ability (K score).

### Predictors of overall high vs. low effort choices

Our primary research question focused on factors that predict the likelihood of choosing high vs low effort/reward options (Table 2; Fig. 4). Here, results of the binomial regression showed that only visual short term memory (K estimate) score significantly predicted whether a participant was in the high effort or the low effort group (*z* = 2.26, *p* = 0.024, *e*^*β*^ = 1.24). Participants with higher working memory ability had a higher probability of being in the high effort group. Depressive traits, chronic stress levels, and reward anticipation did not significantly predict whether participants were in the low vs. high effort groups. Furthermore, a linear model analysis with the same predictors revealed that, once again, only visual short term memory (K estimate) scores significantly predicted the proportion of high effort trials that a participant selected (Table 3).

**Table 2.**
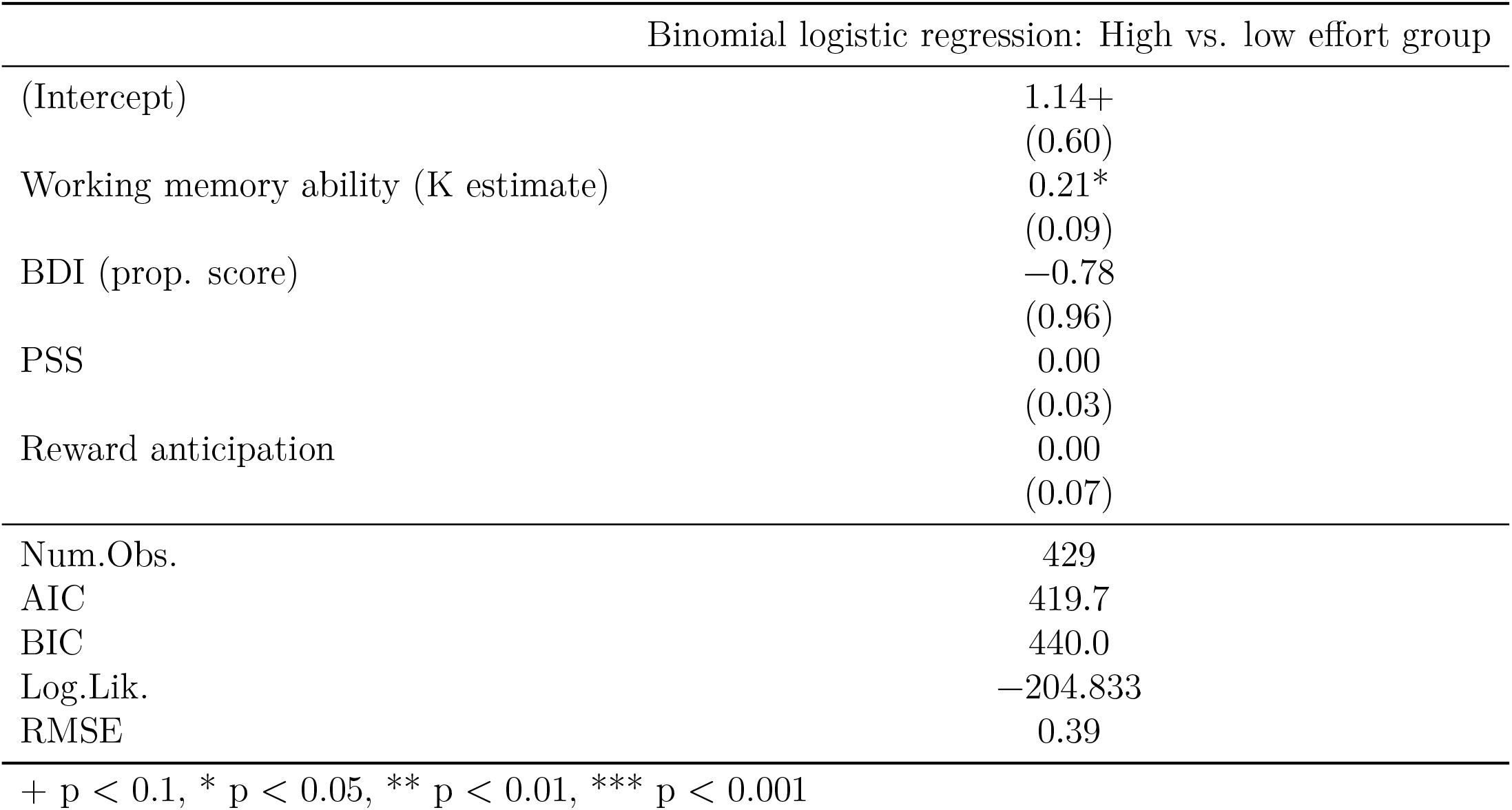
Binomial logistic regression predicting whether participants were in the high effort vs. low effort group. High effort group: > 70% HR trials selected; low effort group: <= 70% HR trials selected. BDI (prop. score) = depression score on the Beck Depression Inventory, PSS = Perceived Stress Scale. Proportion scores are scores divided by total possible score. AIC = Akaike information criterion, BIC = Bayesian information criterion, ICC = intraclass correlation, RMSE = root mean squared error.

**Table 3.** Linear regression of predictors for proportion of high effort trials selected. High effort group: > 70% HR trials selected; low effort group: <= 70% HR trials selected. BDI (prop. score) = depression score on the Beck Depression Inventory, PSS = Perceived Stress Scale. Proportion scores are scores divided by total possible score. AIC = Akaike information criterion, BIC = Bayesian information criterion, ICC = intraclass correlation, RMSE = root mean squared error.

| Linear model: Proportion of high effort trials selected |  |
| --- | --- |
| (Intercept) | 0.84***<br>(0.04) |
| Working memory ability (K estimate) | 0.02**<br>(0.01) |
| BDI (prop. score) | −0.03<br>(0.07) |
| PSS | 0.00<br>(0.00) |
| Reward anticipation | 0.00<br>(0.00) |
| Num.Obs. | 429 |
| R<sup>2</sup> | 0.018 |
| R<sup>2</sup> Adj. | 0.009 |
| AIC | −274.2 |
| BIC | −249.8 |
| Log.Lik. | 143.105 |
| RMSE | 0.17 |

+ p < 0.1, \* p < 0.05, \*\* p < 0.01, \*\*\* p < 0.001

**Figure 4.**
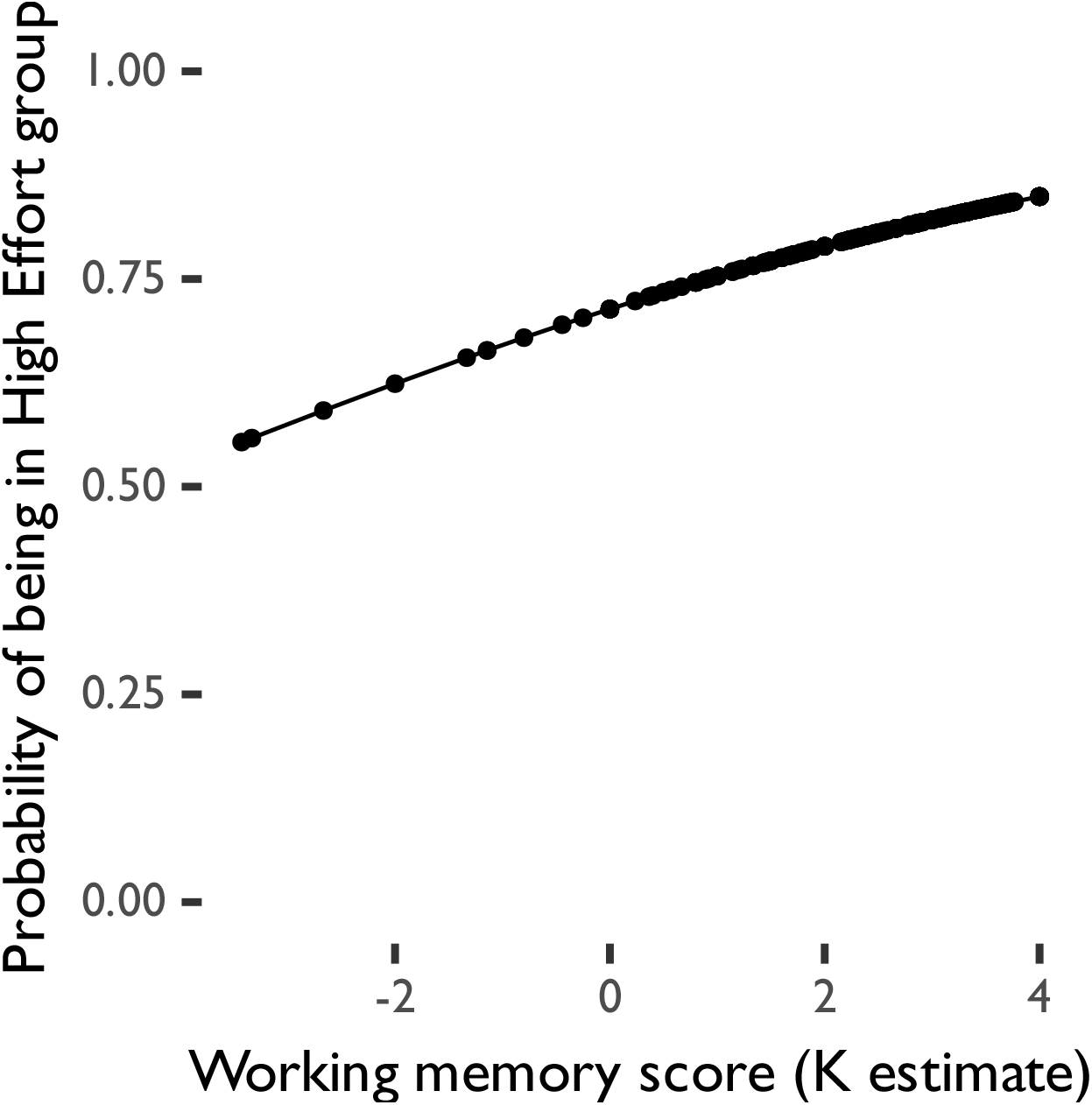
Binomial logistic regression predicting whether participants were in the high effort vs. low effort group. High effort group: > 70% HR trials selected; low effort group: <= 70% HR trials selected.

### Accuracy

In all ANOVAs reported below, all p-values were corrected for multiple comparisons using Greenhouse-Geiser correction. A 2x2 within-between ANOVA (Group x Trial Type; Table 4, Fig. 5A) revealed a main effect of Trial Type on accuracy such that participants showed poorer performance on the more difficult HR trials compared to the easier LR trials (*F*(1, 273) = 724.10, *p* <0.001, *γ* = 0.515). This was qualified by a Trial Type x Group interaction such that participants in the high effort group performed better than those in the low effort group on LR but not HR trials (*F*(1, 273) = 11.81, *p* <0.001, *γ* = 0.017).

**Table 4.** Summary table of accuracy and choice latency by group and effort level chosen. High effort group: > 70% HR trials selected; low effort group: <= 70% HR trials selected.

| Group | Effort level | $M_{Accuracy}$ | $SD_{Accuracy}$ | $M_{Latency}$ | $SD_{Latency}$ |
| --- | --- | --- | --- | --- | --- |
| High effort | high | 0.70 | 0.11 | 1.27 | 1.04 |
| High effort | low | 0.95 | 0.11 | 2.54 | 1.99 |
| Low effort | high | 0.70 | 0.11 | 1.23 | 1.10 |
| Low effort | low | 0.90 | 0.11 | 1.60 | 1.92 |

**Figure 5.**
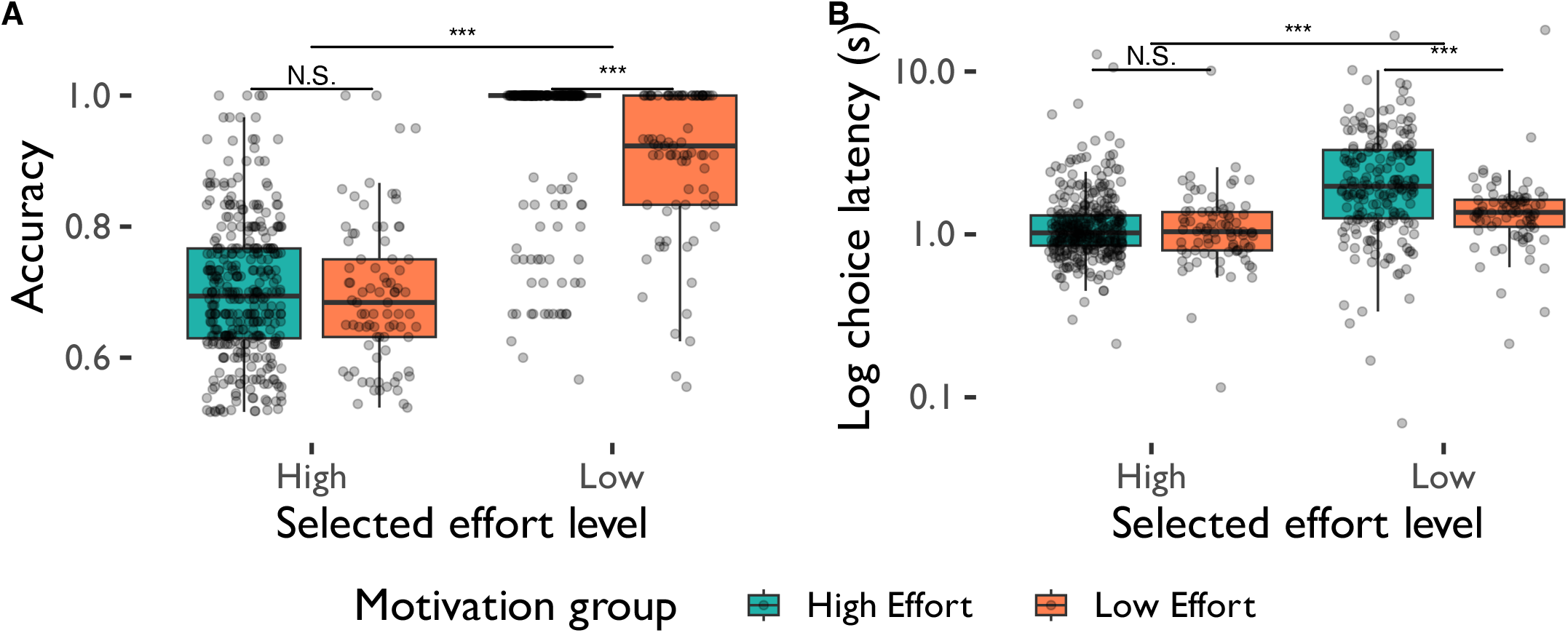
Accuracy and choice latency in the choice phase of the change detection task by motivation group (high effort vs. low effort group). The choice latency plot has been log-transformed on the y-axis to more clearly show small choice latency values. High effort group: > 70% HR trials selected; low effort group: <= 70% HR trials selected.

### Choice latency

A 2x2 within-between ANOVA (Group x Trial Type; Table 4, Fig. 5B) revealed a main effect of Trial Type such that participants were slower to choose HR trials compared to LR trials (*F*(1, 273) = 37.50), *p* <0.001, *γ* = 0.063). Furthermore, high effort participants were slower than low effort participants to make choices across all trial types (*F*(1, 273) = 10.10), *p* = 0.002, *γ* = 0.019). There was also an Trial Type x Group interaction, such that participants in the high effort group took longer to select LR trials than HR trials, while choice latency between high effort and low effort groups did not differ on HR trials (*F*(1, 273) = 37.50), *p* <0.001, *γ* = 0.021).

### Additional predictors of accuracy and choice latency

We additionally evaluated predictors of accuracy on each trial and choice latency for selecting the trial difficulty from individual differences in participant visuospatial short-term memory capacity, mood disorder score, and reward anticipation by sex. A multi-level model analysis revealed that the effort level required in the selected trial significantly predicted accuracy across all trials such that accuracy was higher on low effort (LR) trials compared to high effort (HR) trials. Importantly, of the predictors of interest, only visuospatial short-term memory score predicted acccuracy, such that higher short-term memory ability was associated with higher accuracy (Table 5; Fig. 6A). Trial effort level also predicted choice latency, with slower choices on low relative to high effort trials. Depression scores also predicted choice latency such that higher depression scores predicted faster choices across all trials (Table 6; Fig. 6B) Sex, chronic stress scores, and reward anticipation did not significantly predict accuracy or choice latency.

**Table 5.** Multi-level model analysis coefficients and standard errors for accuracy in the reward phase of the task. BDI (prop. score) = depression score on the Beck Depression Inventory, PSS = Perceived Stress Scale. Proportion scores are scores divided by total possible score. AIC = Akaike information criterion, BIC = Bayesian information criterion, ICC = intraclass correlation, RMSE = root mean squared error.

|  | Multi-level model: Accuracy |
| --- | --- |
| (Intercept) | 0.64***<br>(0.02) |
| Sex | 0.01<br>(0.01) |
| Working memory ability (K estimate) | 0.01**<br>(0.00) |
| BDI (prop. score) | −0.01<br>(0.04) |
| PSS | 0.00<br>(0.00) |
| Reward anticipation | 0.00<br>(0.00) |
| Effort level | 0.25***<br>(0.01) |
| SD (Intercept participant) | 0.04 |
| SD (Observations) | 0.10 |
| Num.Obs. | 542 |
| R2 Marg. | 0.592 |
| R2 Cond. | 0.656 |
| AIC | −840.0 |
| BIC | −801.3 |
| ICC | 0.2 |
| RMSE | 0.09 |

+ p < 0.1, \* p < 0.05, \*\* p < 0.01, \*\*\* p < 0.001

**Table 6.** Multi-level model analysis coefficients and standard errors for choice latency in the reward phase of the task. BDI (prop. score) = depression score on the Beck Depression Inventory, PSS = Perceived Stress Scale. Proportion scores are scores divided by total possible score. AIC = Akaike information criterion, BIC = Bayesian information criterion, ICC = intraclass correlation, RMSE = root mean squared error

| Multi-level model: Choice latency |  |
| --- | --- |
| (Intercept) | 1.25***<br>(0.34) |
| Sex | −0.07<br>(0.17) |
| Working memory ability (K estimate) | 0.05<br>(0.05) |
| BDI (prop. score) | −1.11*<br>(0.54) |
| PSS | 0.01<br>(0.01) |
| Reward anticipation | −0.04<br>(0.04) |
| Effort level | 1.06***<br>(0.13) |
| SD (Intercept participant) | 0.18 |
| SD (Observations) | 1.53 |
| Num.Obs. | 542 |
| R<sup>2</sup> Marg. | 0.114 |
| R<sup>2</sup> Cond. | 0.127 |
| AIC | 2042.9 |
| BIC | 2081.6 |
| ICC | 0.0 |
| RMSE | 1.51 |

+ p < 0.1, \* p < 0.05, \*\* p < 0.01, \*\*\* p < 0.001

**Figure 6.**
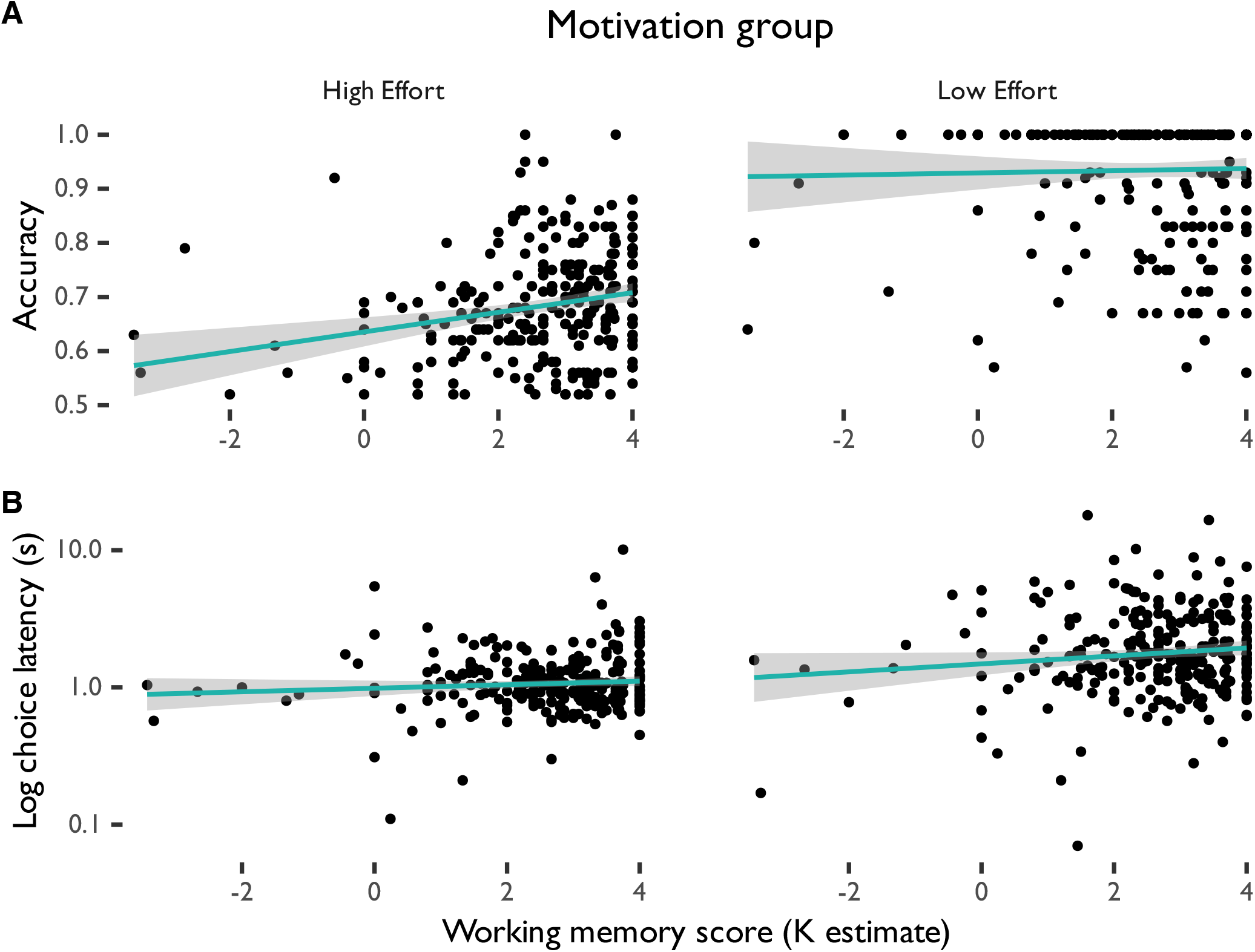
(A) Accuracy and (B) choice latency in the reward phase of the change detection task by working memory ability (K estimate). The choice latency plot has been log-transformed on the y-axis to more clearly show small choice latency values.

### Drift diffusion model

To evaluate group differences in evidence accumulation and bias towards deploying higher amounts of cognitive effort in decision making when choosing HR or LR trials, we fit a hierarchical Bayesian drift diffusion model (Ahn, Haines, and Zhang, 2017) to choice (HR vs. LR trial selected) and corresponding choice latency data on each trial from all participants who had at least one correct response on an HR and an LR trial and who chose at least four trials of each type (*N* = 136, *n*_*Higheffort*_ = 94, *n*_*Loweffort*_ = 42). We then evaluated this overall model fitting data from these participants to explore group-level differences in whether participants selected HR or LR trials more often. For each participant in the high effort and low effort groups, we evaluated four parameters: drift rate (speed of evidence accumulation in deciding on an HR vs. LR trial), starting point (bias towards HR vs. LR trial), boundary separation (extent to which speed and effort choice are traded off between HR and LR trials), and non-decision time (choice latency time unrelated to trial choice, such as time for motor response) (Fig. 7). These parameters capture individual differences in the speed and motivation to make a decision about deploying more or less cognitive effort for reward. After Bonferroni correction for multiple comparisons, we found that high effort group participants had a starting point closer to the HR trial decision boundary (*t*(83.21) = 5.80, *p* <0.001, *d* = 1.05); had wider decision boundaries (*t*(68.49) = 4.38, *p* <0.001, *d* = 0.87) than low effort group participants; and had a significantly higher drift rate (*t*(82.94) = 13.54, *p*<0.001, *d* = 2.46) than low effort group participants. However, participants did not significantly differ in non-decision time (*t*(105.65) = 2.34, *p* = 0.192, *d* = 0.39). These findings suggest that participants who were in the high effort group required more evidence to select low but not high effort trials, were more strongly biased towards selecting high effort/high reward trials, and were quicker to accumulate evidence for these choices, than those in the low effort group.

**Figure 7.**
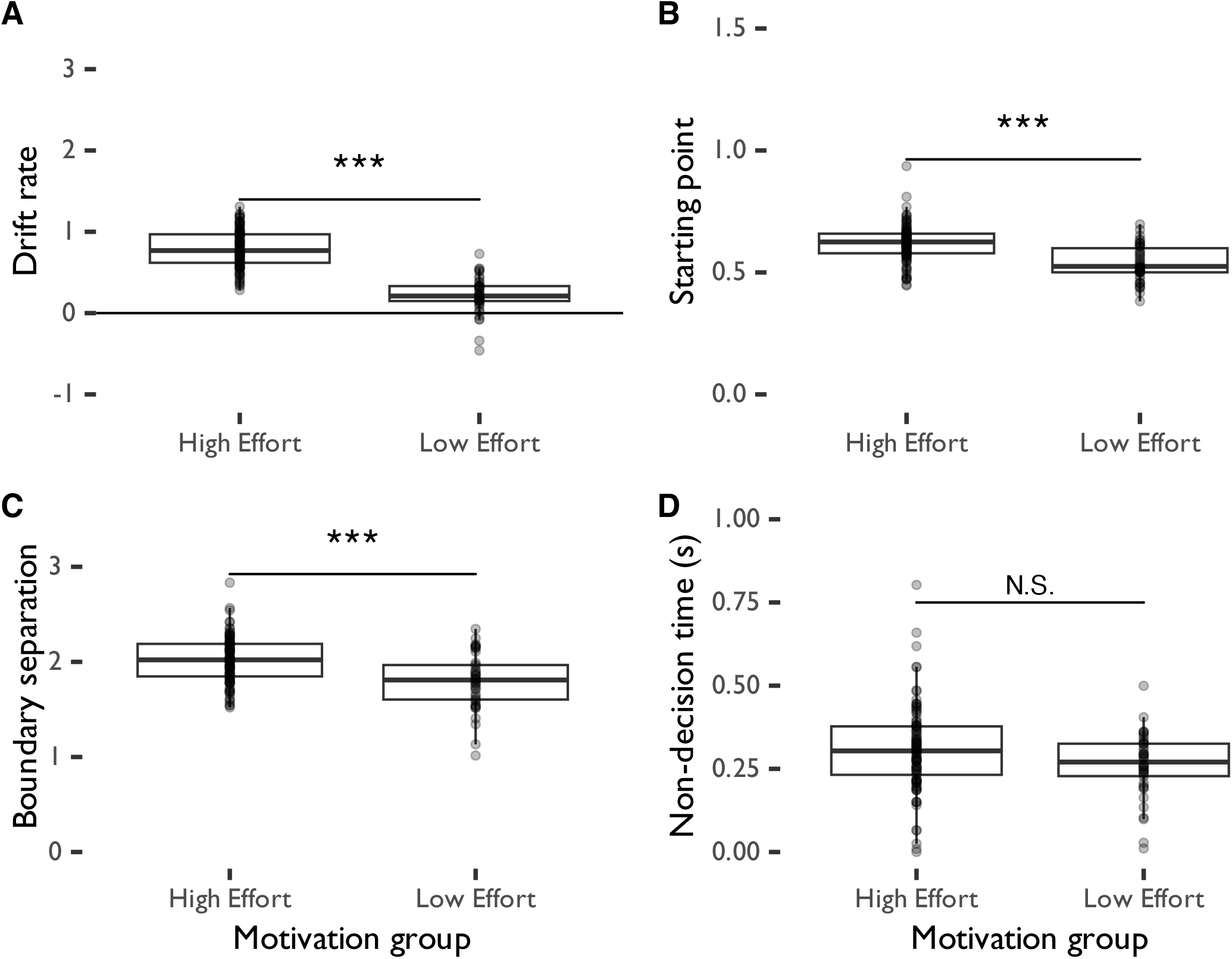
Outputs of a hierarchical Bayesian drift diffusion model fit to all participants in the effort choice task, divided by effort group. High effort participants had a significantly higher (B) starting point and (C) boundary separation compared to low-effort participants, but did not significantly differ on (A) drift rate or (D) non-decision time compared to low-effort participants.

## Discussion

In this study, we investigated whether working memory performance, depressive traits, and chronic stress levels influence people’s motivation and ability to consistently deploy cognitive effort for reward. Results showed that greater working memory ability significantly predicted the likelihood of systematically choosing high effort/high reward trials (whether the participant was classified in the high vs. low effort group), whereas chronic stress, depression trait levels, and reward anticipation were not predictive of effort deployment choices. Additionally, participants required more evidence to select LR trials compared to HR trials. Drift diffusion modeling further indicated that those who were categorized as high-effort participants required less evidence to select high effort/high reward trials, and were more strongly biased towards selecting these trials, than low-effort participants. Furthermore, working memory ability significantly predicted accuracy, while depressive traits predicted choice latency; both factors significantly differed depending on the effort required in the trial.

These results partially recapitulate findings from rodent research using the rCET regarding differences in performance and decision-making biases between those who select high effort trials more often, and those who select them less often. As in Cocker, Hosking, Benoit, and Winstanley (2012), participants who chose high effort trials over 70% of the time performed better than those who chose high effort trials 70% or less of the time, on LR - although not HR - trials. Continually deploying high amounts of cognitive effort could be fatiguing; the high effort group’s lower performance on HR trials could reflect the ineffectiveness of this high effort choice strategy.

High effort group participants, who generally selected HR rewards more often and who were more biased towards selecting HR trials than low effort group participants, appear to have spent more time deliberating before selecting a LR trial than low effort group participants. Choice latency indicates how long participants spend when deciding which effort level to choose. Thus, they may have been trading off the benefits and costs of selecting a LR trial more consciously than low effort group participants were (Pruessner, Barnow, Holt, Joormann, and Schulze, 2020; Sandra and Otto, 2018), even as they performed significantly better than low effort group participants on LR but not HR trials. These tradeoffs could shift over time in the task as participants become fatigued or bored with the task, or found a new effort deployment strategy that better optimized rewards or task enjoyment.

Furthermore, our findings align with past work in rodents (Hales, Silveira, Calderhead, Mortazavi, Hathaway, and Winstanley, 2024) in that our drift diffusion model results showed wider decision boundaries between HR and LR trials for high effort group compared to low effort group participants, as well as a starting point closer to HR trials for high effort group participants. Given that the decision available on each trial is the same, individual differences in this initial level of bias towards selecting high effort trials - together with random noise - is what drives differences in trial difficulty choice between participants as modeled by the drift diffusion model. Therefore, the drift diffusion model may not be capturing meaningful, systematic patterns of effort deployment choices. However, it still provides an exploratory evaluation of the underlying cognitive processes that could be driving effort deployment choices. Additionally, as with rodents, participants in the low effort group made faster choices than those in the high effort group on LR trials. Furthermore, high effort group participants showed steeper drift rates than low effort group participants - while rodents in the equivalent high effort group had steeper drift rates when they subsequently made correct responses for their given choice (Hales, Silveira, Calderhead, Mortazavi, Hathaway, and Winstanley, 2024). Taken together, these findings could suggest similarities between humans and rodents in the influence of cognitive capabilities on performance when deploying more cognitive effort, as well as similarities in biases towards deploying more effort or wanting higher rewards. Furthermore, given the conditionality of the drift rate findings in rodents, strategies driving impulsiveness or choice motivations to select a HR or LR trial could differ between humans and rodents. This could be explained by differences in the strength of reinforcement provided by the reward, which was a secondary reinforcement (points towards a monetary reward) in our task but a primary reinforcement (sugar pellets) in the rCET (Cocker, Hosking, Benoit, and Winstanley, 2012; Hales, Silveira, Calderhead, Mortazavi, Hathaway, and Winstanley, 2024), or by other differences between our task and the rCET as well as between humans and rodents in terms of how reward motivation is processed. These differences were not driven by reward anticipation, which did not differ between high and low effort groups (*t*(122.95) = 0.17, *p* = 1, *d* = 0.02).

Additionally, previous studies with the rCET have shown no overall accuracy differences between high and low effort group participants across either LR or HR trials (Hosking, Cocker, and Winstanley, 2016; Silveira, Wittekindt, Mortazavi, Hathaway, and Winstanley, 2020). However, in a recent, larger analysis, Hales, Silveira, Calderhead, Mortazavi, Hathaway, and Winstanley (2024) showed that rodents choosing more HR trials were slightly more accurate on HR than LR trials when completing the trials themselves, while rodents choosing more LR trials were slightly more accurate on LR than HR trials when completing these trials. Although participants practiced task contingencies ahead of the main choice phase and the task difficulty was calibrated to working memory ability, individual differences in working memory - which predicted whether a participant selected high or low effort trials more often - still predicted these performance differences in humans. Furthermore, chronic stress - which can negatively impact availability of executive resources - did not explain accuracy or choice latency in this task. Thus, within the constraints of a binary effort choice task, participants’ willingness to deploy cognitive effort was explained more by individual differences in executive function (working memory capability) (Sandra and Otto, 2018) rather than affective factors or the effects of affective factors on executive function (Slattery and Cryan, 2017). Note that although we did not measure executive control as a global construct in this study, visuospatial working memory ability is typically described as a process that falls withing the suite of executive functions (Miyake and Friedman, 2012).

Our drift diffusion model results also suggest that high effort group participants had to overcome a bias towards HR trials in order to select LR trials. This process could be driven by adjustments to effort deployment strategies based on performance, as suggested by recent interpretations of the Expected Value of Control theory (Shenhav, Prater Fahey, and Grahek, 2021). Through this framework, choosing a high proportion of high effort trials could be seen as optimizing effort deployment and making considered, model-based, decisions based on performance - as reflected in increased choice latency among high effort group participants for LR trials - whereas selecting a high proportion of low effort trials could entail more model-free, rapid, and random decisions. We conducted a follow-up linear regression analysis to evaluate whether performance on the previous trial predicts performance on the current trial as a function of effort motivation group and trial type. Performance on the prior trial, but not effort motivation group, significantly predicted performance on the current trial (*t*(68955) = 8.45, *p* <0.001), suggesting that, overall, participants maintained a constant performance-based strategy that was not explained by effort choice decisions. Furthermore, participants with higher levels of depressive traits responded significantly faster across all trials, a result that runs counter to past work suggesting increased choice latency in reward tasks in rodents (Hales, Stuart, Griffiths, Bartlett, Arban, Hengerer, and Robinson, 2023) or humans (Di Schiena, Luminet, Chang, and Philippot, 2013) with depressive-like traits. However, after Bonferroni correction, depressive traits and choice latency were not significantly correlated across all participants and trials (*t*(702) = -1.21, *p* = 1, *r* = -0.05). As depressive traits did not impact accuracy, these findings could instead suggest a lack of engagement with decision-making in the task characterized by higher levels of automatic responding given high levels of depressive traits (Teachman, Joormann, Steinman, and Gotlib, 2012).

Our findings suggest that choices to deploy cognitive effort for reward are primarily driven by participants’ cognitive capacity (as measured by visuospatial working memory ability) and not by depressive traits, chronic stress, or reward anticipation, in participants within a non-clinical range of depression traits and chronic stress. Potentially, the aversiveness of expending cognitive effort for participants with lower working memory capacity may have overridden overall sensitivity for reward. Additionally, although our population exhibited a wide range of depression scores - including those above clinical thresholds (Wang and Gorenstein, 2013) - this population was non-clinical. In contrast, much work on the impact of reward anticipation on cognitive flexibility and related constructs has focused on participants who were clinically diagnosed with depression (Alloy, Olino, Freed, and Nusslock, 2016; Terpstra, Vila-Rodriguez, LeMoult, Chakrabarty, Nair, Humaira, Gregory, and Todd, 2023) or on rodents in which chronic stress was directly induced (Watt, Weber, Davies, and Forster, 2017). In order to obtain the highest accuracy and reward possible, participants may have titrated the task’s difficulty - through choosing a higher proportion of LR trials - to a level at which they can consistently complete the task and at which the value of their effort was greatest (Shenhav, Botvinick, and Cohen, 2013).

We found no significant sex differences in performance or choice latency in our study. Women generally present more with depression than men (Parker and Brotchie, 2010) and are also more impacted by chronic stress across the lifespan (Hodes and Epperson, 2019). However, the results of our study were primarily driven by visuospatial short-term memory ability, where sex differences are overall smaller and women may be better than men at memory for location (Voyer, Voyer, and Saint-Aubin, 2017) - a relevant measure in our task. Our undergraduate psychology student sample strongly skewed female; in future studies, it would be important to obtain a more balanced sample in order to further explore potential sex differences.

There are a number of caveats we must consider when interpreting these results. First, the study was conducted fully online on participants’ own devices. While this allowed us to obtain a larger and better powered sample, differences in screen size and background distractions could have impacted participants’ ability to encode the positions and colours shapes on each trial. These differences may have made the task more difficult than originally intended for some participants. However, participant performance was still near ceiling on LR trials and very high on HR trials, so this was unlikely to have a large effect.

Second, high effort trials never gave low rewards, or vice versa. This may impact our ability to evaluate whether it was sensitivity to effort or to reward that more strongly drove participant choices in the task. However, reward anticipation - as measured in a task that involved accumulating did not predict choices for high effort deployment. This suggests that effort costs could be weighed more strongly than rewards by participants when making effort choices.

Third, the set sizes used on the change detection task were modified from those originally used in Luck and Vogel (1997). The set size of 10 was removed from the pool of trials as pilot participants had very low performance at this set size. Although this change reduced the possible variety of trials used, participants still reported a maximum set size of 8 to be highly challenging. Furthermore, the choice structure of the task incorporated only two of the set sizes. This could have made the task excessively easy for some participants - as reflected in the ceiling effect on LR trials - and excessively difficult to others. However, we did titrate task difficulty according to participants’ visual short term memory ability, reducing the likelihood of the task difficulty not matching participant ability. Future studies could use a finer-grained, parametric design that offers choices at more difficulty levels to examine whether motivation continuously varies with effort levels (Sayalı, Heling, and Cools, 2023).

Fourth, the method used to calculate the K estimate slightly differed between the task - used to set the criterion level - and our analysis - used to establish the K estimate for each participant. In the task, the K estimate was calculated based on the current hit and false alarm rate after every trial, with the final K estimate being used to set the criterion for the choice phase of the task. In our analysis, we calculated the K estimate based on the full hit and false alarm rate at each set size. Because of rounding differences in these calculations, the K estimate values used in our analysis differ from those in our task by *M* = 0.01, *SD* = 0.63. We consider this difference to be small enough as to not have an effect on the difficulty of the task.

Fifth, participants were told that the points they received would be proportional to the monetary reward received, which could motivate them towards higher effort choices regardless of their effort capabilities - shifting their effort-reward tradeoffs. However, such a strong effort incentive was necessary to motivate participant performance given the difficulty of the high effort trials for many participants. Participants may have also individually differed in compliance with reading instructions or in their motivation for reward. Such factors could be investigated in a follow-up study incorporating qualitative evaluations of participant experiences of the task.

Sixth, differences in performance that are explained by the K estimate could also be explained by the differences in set sizes available associated with being above or below the K criterion of 3 on set size = 4. However, the set sizes at each level of criterion were established with piloting such that the difficulty of each criterion level was matched as closely as possible. Last, although our task only had a single probe whose colour had to be compared to the original probe, participants had to attend to the whole display to compare the single probe to the existing square at any given location on screen. As such, we used the whole-display K estimate measure to calibrate participant performance in our task (Rouder, Morey, Morey, and Cowan, 2011). A single-probe K estimate measure reflects lower effort demands than the whole-display measure, as the whole-display measure requires the status of all shapes on screen to be held in memory for the probe – whereas the single-probe measure only involves evaluating one shape in one position. However, as the position of the probe shape in our task is randomized, participants must still attend to the whole display to correctly identify any changes in the probe’s colour. As such, the whole-display K estimate measure is more relevant for our task’s structure and cognitive effort demands, and the results could differ if we use the single-probe measure.

Future studies could make further use of information about effort deployment choices that can only be captured in human studies. As many human cognitive effort tasks - unlike rodent tasks - do not require substantial training, a future study could examine effort choices given a wider range of effort and reward levels, as well as test how preference for effort changes given high effort trials with low reward and vice versa. Such a task could more richly capture individual differences in human cognitive effort deployment decisions. Furthermore, an expanding field of research uses qualitative approaches to investigate why participants make the judgments they do about effort deployment, as well as ask about participant experiences of task difficulty and the value of rewards vs. the effort required to obtain them (Vásquez-Rosati, Montefusco-Siegmund, López, and Cosmelli, 2019). Understanding the phenomenology of cognitive effort deployment choices would help us extend our understanding of how and why we make decisions to deploy more or less cognitive effort for reward, beyond traditional self-report measures. Participant descriptions of their experiences in the task could indicate whether boredom and fatigue, in addition to reward motivation, impacted effort deployment choices. This approach would also allow us to further explore how rodent-based constructs of cognitive effort deployment are reflected in human decision-making, while extending our understanding of these constructs with human-specific experiential information. Additionally, exploring how the brain represents information about effort task difficulty - especially in regions relevant to cognitive control like the dorsal anterior cingulate cortex (Shenhav, Cohen, and Botvinick, 2016; Yee, Crawford, Lamichhane, and Braver, 2021) - through fMRI would help us better understand how motivational states in cognitive effort decision-making are reflected in neural circuitry.

Our study suggests that, as in rodents, significant individual differences exist in the human tendency to choose harder but more rewarding options in cognitive effort tasks. Using a visuospatial working memory task, we show that individual differences in accuracy on the trial and choice latency for the trial type chosen are primarily driven not by depressive traits or chronic stress - as has previously been shown in the rodent literature - but instead by working memory capability. These findings may help to inform clinical interventions aimed at increasing motivation to seek rewards and engage in work in everyday life, and illustrate the importance of translating and extending rodent work with human-specific measures.

## Data and code availability

All task code, stimuli, data and code used to generate this manuscript and the figures are available at https://osf.io/c4h7s/.

